# Unraveling the regulatory role of miRNAs responsible for proanthocyanidin biosynthesis in the underutilized legume *Psophocarpus tetragonolobus* (L.) DC

**DOI:** 10.1101/2021.07.24.453638

**Authors:** Sagar Prasad Nayak, Priti Prasad, Vinayak Singh, Abhinandan Mani Tripathi, Sumit Kumar Bag, Chandra Sekhar Mohanty

## Abstract

The underutilized legume winged bean (*Psophocarpus tetragonolobus* (L.) DC.) is deposited with various degrees of proanathocyanidin (PA) or condensed tannin (CT) on its seed-coat. PA content of two different lines of *P. tetragonolobus* was estimated and accordingly they were denoted as high-proanthocyanidin containing winged bean (HPW) and low-proanthocyanidin containing winged bean (LPW). The level of PA-content varied as 59.23 mg/g in HPW and 8.68 mg/g in LPW when estimated through vanillin-HCl assay. The identification and quantification of catechin and epigallocatechin gallate were estimated in a range of 63.8 mg/g and 2.3mg/g respectively in HPW whereas only epigallocatechin gallate was reported in LPW line with a value of 3 mg/g. A comparative miRNA profiling of the leaf-tissues of these contrasting lines of *P. tetragonolobus* revealed a total of 139 mature miRNAs. Isoforms of known novel miRNAs were also identified in this study. Differentially expressed miRNAs e.g., miR156, miR396, miR4414b, miR4416c, miR894, miR2111 and miR5139 were validated through qRT-PCR analysis. Target prediction of the identified miRNAs especially miR156, miR396, miR4416b shows that they have a potential role in the proanthocyanidin biosynthesis of *P. tetragonolobus*. The study will provide the basis for understanding the role of miRNAs in regulating the biosynthesis of proanthocyanidin.

## Introduction

Flavonoids are the largest and widely distributed group of secondary metabolites in the plant-kingdom (Winkel-Shirley, 2001). In addition to provide color, they confer protection to the plant. They take part in plant growth, development, transport, signaling and many other vital activities (Koes et al., 2005). More than 6000 different groups of flavonoids with diverse biological functions have been reported till date (Falcone Ferreyra et al., 2012). Though they are easily detectable in flowers as pigments, they are widespread in occurrence and are found in several parts of the plant. They are found across the plant kingdom and through plant-based foods such as fruits, vegetables and beverages they enter into the food chain (Dewick, 2009). Flavonoids have been divided into various sub-groups like flavones, flavanols, isoflavones, flavanones and anthocyanins (Panche et al., 2016).

Proanthocyanidins (PAs) or condensed tannins (CTs) are the oligomeric flavonoids that contribute significantly to the dietary polyphenols in the plants (Santos-Buelga and Scalbert, 2000). Almost all parts of the plant including leaves synthesize PA. However, plants deposit PA preferably on the outer integument of the seeds (Xu et al., 2014). Despite their wide range of occurrence in different plant parts, they are considered as anti-nutrient because of their interacting-property with proteins (Duodu and Dowell, 2019). They form complexes with food proteins and lower the feed-efficiency (Chen et al., 2017; Reddy et al., 1985). As they possess multiple functional groups, so they can easily make bonds with protein and carbohydrate molecules (Fraga-Corral et al., 2020). Apart from being an anti-nutrient, proanthocyanidins also provide beneficial health effects by mitigating inflammation and oxidative stress (Beecher, 2004) for which it has gained pharmaceutical attentions nowadays. PA biosynthesis takes place through phenylpropanoid pathway by sequential action of both early and late biosynthetic genes (Rauf et al., 2019). Involvement of an array of enzymes, proteins along with the corresponding transcriptional regulators have made this pathway more complex to understand. The synthesis of proanthocyanidin requires the action of a ternary complex of three different classes of transcription factors viz. R2R3-MYB, bHLH and WD40 (Li et al., 2018; Li, 2014). These transcription factor complexes activate the expression of late biosynthetic genes catalyzing the synthesis of downstream compounds. Genes specific to PA biosynthesis are leucoanthocyanidin reductase (LAR) (Abrahams et al., 2003), anthocyanidin synthase (ANS) or leucoanthocyanidin dioxygenase (LDOX) and anthocyanidin reductase (ANR). LAR uses leucocyanidin as a substrate to produce catechin - the monomeric unit of proanthocyandin, which further condenses to bioactive PA. In this process, ANS plays a significant role in the production of colored anthocyanin (He et al., 2008). ANS oxidizes leucoanthocyanidin to anthocyanidin-precursor molecule for the biosynthesis of anthocyanin. The remaining anthocyanidins get converted into epicatechin (monomeric units of PA) through the action of anthocyanidin reductase. This is coded by the BANYULS gene (Xie et al., 2003). The final condensing enzymes for catalyzing the polymerization step of PA production are still unknown and the process of condensation is believed to occur through oxidation. The monomeric units of proanthocyanidin viz. catechin and epicatechin are synthesized on the cytosolic face of endoplasmic reticulum (Brillouet et al., 2013), through the action of a multi-enzyme complex (Saslowsky and Winkel-Shirley, 2001) and the polymerization takes place inside the chloroplast derived organelle called “tannosome” (Brillouet et al., 2013). In some plants, the catechin and epicatechin molecules are galloylated to form the PA, they are linked to many health benefits (Xu et al., 2016).

MicroRNAs (miRNAs) are small regulatory RNAs that regulate almost all aspects of plant-growth and development (Sunkar et al., 2012). The role of miRNAs is well-studied from cellular-life to stress responses in plants (Alptekin et al., 2016). High-throughput sequencing and computational analysis facilitated the understanding of miRNAs, especially their biogenesis, evolution, potential targets and regulatory effect on gene expression (Gupta et al., 2017; Luo et al., 2013). Mature miRNA sequences are of (19-24) nucleotides (Bartel, 2009) in length having functional role in post-transcriptional gene silencing (Baumberger and Baulcombe, 2005). It preferably targets the transcription factors (TFs) (Rubio-Somoza and Weigel, 2011; Sreekumar and Soniya, 2017) like miR156 targets TF SQUAMOSA PROMOTER BINDING-LIKE (SPL9) and miR172 targets APETALA2 in *Arabidopsis* (Jangra et al., 2018). Efficient identification of miRNAs in large number of plant species and elucidation of their functional role in regulating different biological processes in different life-stages of plants have been performed by many research groups. Efforts have also been made towards identifying miRNAs involved in phenylpropanoid biosynthetic pathway. In *Canna*, five miRNA families have been identified to play possible roles in phenylpropanoid biosynthetic pathway (Roy et al., 2016). It has been reported that, miR858a targets R2R3-MYB transcription factors to regulate flavonoid biosynthesis in *Arabidopsis* (Sharma et al., 2016) and miR156 targets squamosa promoter binding like (SPL) proteins for accumulation of anthocyanin and enhanced levels of flavonols (Gou et al., 2011). Differential expression of miRNAs in leaf and flower tissues with relations to their flavonoid levels have been checked in *Osmanthus*. MiRNAs with probable role in flavonoid biosynthesis were detected (Shi et al., 2021). Moreover, genes associated with glycosylation and insolubilisation of tannin precursors are the possible targets of dka-miR396g and miR2911 in persimmon fruit (Luo et al., 2015)

Winged bean (*Psophocarpus tetragonolobus* (L.) DC.), is a crop that stands in the category of underutilized legume despite being a good source of protein and oil (Singh et al., 2017). The PA content in winged bean varies significantly among the *P.tetragonolobus* cultivars (0.3 to 7.5 mg/g) (Tan et al., 1983). *P. tetragonolobus* was reported to possess soybean equivalent nutrients (Mohanty et al., 2013) Presence of PA in *P.tetragonolobus* is probably one of the reasons for its underutilization as it limits the bioavailability of nutrients. Owing to their involvement in various biological processes, miRNAs have been identified, analyzed and experimentally validated in many leguminous plants including *Phaseolus vulgaris* (Pelaez et al., 2012), *Cajanus cajan* (Kompelli et al., 2015; Nithin et al., 2017), *Vigna unguiculata* (Gul et al., 2017)*, Caragana intermedia* (Zhu et al., 2013) and *Cicer arietinum* (Hu et al., 2013). But, till date, there is no report on miRNAs and their associated functions in *P. teragonolobus*.

To systemically identify the miRNAs regulating PA biosynthesis in *P. tetragonolobus,* small RNA sequencing was carried out in the leaf tissues of two cultivars with contrasting levels of proanthocyanidin content namely HPW (high proanthocyanidin containing winged bean) and LPW (low proanthocyanidin containing winged bean). In this study, conserved and novel differentially expressed miRNAs between the contrasting lines along with their putative targets were identified. The significant novel miRNAs in *P. tetragonolobus* along with their secondary structures were predicted. Expression profiling of conserved and novel miRNAs were also investigated through the qRT-PCR. This study provided an insight into regulatory network on PA metabolism in winged bean. The proposed network of miRNA-based regulation of PA biosynthesis and the generated data shall be helpful in future study and improvement program of *P. tetragonolobus*.

## Materials and methods

### Estimation of proanthocyanidin

Quantification of proanthocyanidin content was carried out through vanillin-HCl assay (Price et al., 1978). Approximately, 200 mg of *P. teragonolobus* leaves at an early stage of lignification was collected and weighed for extraction with 10 ml of methanol. The collected supernatant was further processed for spectrophotometric analysis at 500 nm with catechin equivalent standard (CES) (Price et al., 1978). Based on the analysis and screening, diverse lines with high proanthocynidin content (HPW) and low-proanthocyanidin content (LPW) of *P. tetragonolobus* were identified for further analysis.

### Sample preparation, detection and quantification of PA units

Leaves from the field-grown plants were collected and extracted with methanol at concentration of 1 mg ml^-1^. The supernatant was collected and filtered for HPLC analysis. For preparation of standards, catechin and epigallocatechin gallate were weighed and dissolved in methanol and diluted to the required concentrations. The gradient mobile phase consisting of component A (acetonitrile) and component B (water) was used. The elution of mobile phase gradient program was as (0−13) min, 21% B; (13−38) min, 36% B; (38−50) min, 50% B; (50−60) min, 21% B. Constant flow at 1 ml min^-1^ was maintained and the investigated compounds were determined at 254 nm. The standard and sample injection volume were 20 μl. Each compound was identified with the help of retention time and by spiking with the standards under the same conditions.

### RNA isolation, library preparation for sequencing

HPW and LPW lines of *P. teragonolobus* were grown and maintained in the garden of CSIR-National Botanical Research Institute, India. Fresh leaf samples were frozen in liquid nitrogen and total RNA was isolated using mirVana^TM^ miRNA isolation kit (Thermo Fisher Scientific, USA). The NEB next small RNA sample preparation protocol was used to prepare the sample sequencing library. Illumina adapters in the kit were directly, and specifically, ligated to miRNAs. The libraries were prepared as per the manufacturer’s protocol. RNA 3’ adapter was specifically modified to target miRNAs and other small RNAs that have a 3’-hydroxyl group resulting from enzymatic cleavage by dicer or other RNA processing enzymes. The adapters were ligated to each end of the RNA molecule and the reaction was performed to create single stranded cDNA. The cDNA was then PCR amplified using a common primer and a primer containing one of the 48 index sequences. After library preparation, they were sequenced on Illumina HiSeq 2500 platform with 1×50bp read lengths.

### MiRNA data processing and identification of conserved miRNAs

Raw reads of 50 bp from the HPW and LPW sequenced library were subjected for the removal of the adapter sequences through the srna workbench toolkit (Stocks et al., 2012). The filtered reads with a range of 16 to 36 bp length were subjected to blast with the RFAM database v13 (Kalvari et al., 2018). This process helped us to remove the small RNAs other than the miRNAs from both the library, separately. The reads that showed high similarity with full coverage were excluded and assigned as rRNA, sRNA, tRNA and snoRNA. Unmapped sequences from the RFAM database were further aligned against to the miRBase v21 (Kozomara et al., 2019) by mirProf pipeline to identify the known conserved and non-conserved miRNAs. Default settings were used with the depth of minimum 3 reads and 2 mismatches for miRNAs identification. *Glycine max* was selected as the reference genome for identification of conserved and known miRNAs, as the genomic information of *P. teragonolobus* is not available in the public domain database till date.

### Prediction of novel miRNAs and isoforms

Novel miRNAs prediction in the sequenced library was carried out through mirDeep2 tool (Friedlander et al., 2012). Unmapped reads in both the libraries, were pooled and mapped on *Glycine max* reference genome by bowtie v1 algorithm (Langmead and Salzberg, 2012). The sequences of 18 to 25 nucleotide length were selected for defining the novel miRNAs in the datasets by allowing one mismatch in the seed region with significant randfold *p*-value. Randfold calculated the minimum folding energy for generating the stable secondary structure.

The isoforms of the conserved miRNAs were also deduced through custom scripts. Unmapped miRNA reads that showed sequence similarity with *Glycine max* miRNAs with 1 to 7 nt overhangs, were classified as the isoforms of conserved miRNAs. These overhangs present either at 5’ or 3’ ends in the pooled datasets.

### Quantification and differential expression analysis of conserved miRNAs

Identified, conserved miRNAs of HPW and LPW lines of *P. tetragnolobus* were quantified for the differential expression analysis using DESeq package in R (Anders and Huber, 2010). Raw mapped counts of identified conserved miRNAs were used as the input for calculation of differentially expressed miRNAs with FDR value ≤ 0.05 and log2 (fold change) > 1. To understand the functions of known/conserved and differentially expressed miRNAs in *P.tetragonolobus*, their putative targets were predicted in psRNATarget server, a tool for plant smallRNA target prediction (Dai et al., 2018). The default parameters for miRNAs target prediction were used by considering *Glycine max* as a reference genome.

### Functional enrichment of conserved and novel miRNAs

The putative targets of differentially expressed miRNAs between HPW and LPW lines of *P. tetragonolobus* were visualized through cytoscape (Shannon et al., 2003). Significant Gene Ontology (GO) enrichment of the identified conserved and novel miRNAs were carried out through Agri GO software v2 (Tian et al., 2017) with the Singular Enrichment Analysis (SEA) tool and visualized through ggplot2 library in R-package. The involvement of identified miRNAs in different secondary metabolic pathways were classified separately based on mapman pathways database(Thimm et al., 2004).

### Validation of miRNAs through qRT-PCR

For the real time expression and validation in *P. tetragonolobus*, twelve miRNAs (known and novel) which were expressed in both the libraries were randomly selected. The corresponding primers were designed through Primer blast and synthesized from GCC India Private limited (Supplementary Table 3). A total of 500 ng of RNA were isolated from the leaves of two diverse lines (HPW and LPW) of *P. tetragonolobus* using SuperScript III Reverse Transcriptase (Invitrogen, USA). Stem loop cDNAs were synthesized for each miRNA, individually. The expression levels of all the selected miRNAs were quantified by real time PCR using Applied Biosystems 7500 Fast Real-Time PCR System. Each reaction mixture contained a total volume of 20 µl with 2 µl cDNA, 2 µl of primers (forward and reverse), 10 µl Syber Green PCR Master mix (Thermo fisher Scientific, USA) and 6 µl nuclease free water. Reactions were carried out using three replicates for each sample. For normalization, the average Ct value of multiple genes were obtained to get the value of ΔCt (Sun et al., 2015). The fold change value was calculated by 2^−ΔΔCT^ method. U6 (RNU6-1) snRNA was chosen as an internal control for the expression analysis.

## Results

### Estimation of PA and their structural units

The PA content of the leaves of contrasting lines of proanthocyanidin-containing *P. tetragonolobus* (HPW and LPW) was estimated through modified vanillin-HCl assay (Price et al., 1978). On quantification, the PA content of HPW line of *P. tetragonolobus* was reported to be 59.23 mg/g while LPW had 8.68 mg/g in the leaf tissues (**Figure 1A)**. The PA content of these two lines displayed a significant difference. Identification and quantification of different monomeric units of PA (i.e., catechin and epigallocatechin gallate) in the leaf tissues of HPW and LPW lines of *P. tetragonolobus* was carried out on HPLC platform for comparative biochemical analysis **(Fig S1)**. The identified molecules were calibrated at 280 nm and their levels were quantified in both the lines (**Figure 1B**). The presence of catechin and epigallocatechin gallate was reported in HPW line with a value of 63.8 mg/g and 2.3mg/g respectively. The LPW line reported the presence of epigallocatechin gallate in the leaf tissues with a value of 3 mg/g.

**Fig. 1.**
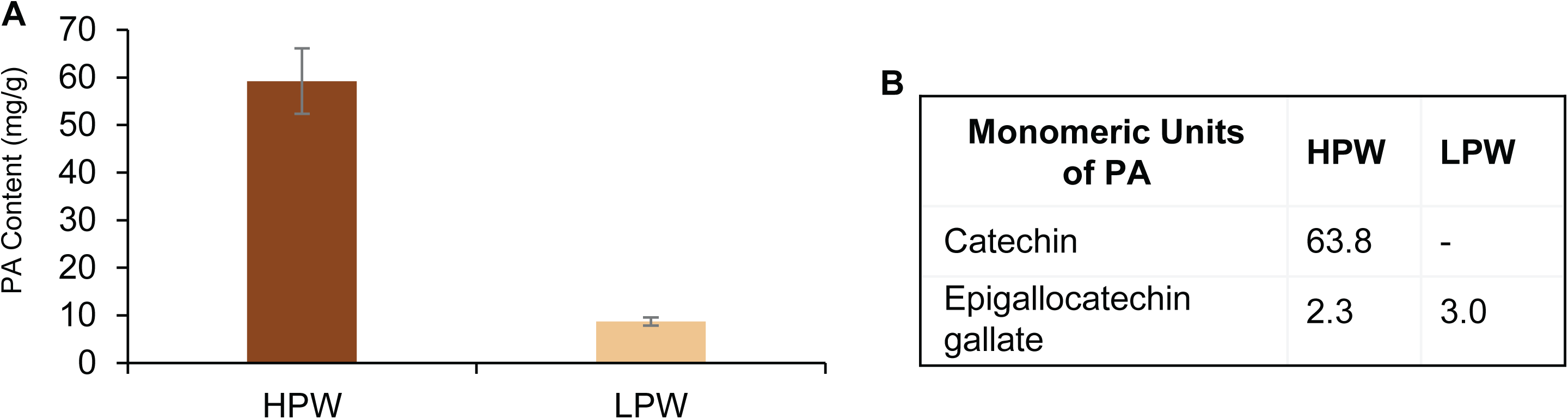
**A.** Proanthocyanidin (PA) composition in leaf tissues of two selected lines of *P. tetragonolobus*. PA content was measured in mg/g of leaf tissues. Higher PA content line was referred as a High Proanthocyanidin Winged bean containing Line (HPW) wheareas Low PA content line was referred as a Low Proanthocyanidin Winged bean containing Line (LPW) **B.** Qualitative and quantitative analysis of monomeric units of proanthocyanidin in methanolic extracts of leaf tissues in two contrasting lines (HPW and LPW) of *P. tetragonolobus* using HPLC method.

### Small RNA library preparation and data processing

Small RNA sequencing of HPW and LPW lines of *P.tetragonolobus* was conducted through Illumina HiSeq 2500 platform that generated an average of ∼50 million single-end reads of 50bp length. The generated data is submitted in the public domain database of NCBI with the SRA accession number SRR15115657 to SRR15115660. After the removal of the low-quality reads, adaptor contaminations and redundancies, approximately five and twelve million unique filtered reads of 16 to 35 nt lengths were selected in HPW and LPW libraries, respectively (**Table 1**). The clean reads of both the libraries were mapped to RFAM database to filter out any contaminated small RNA other than miRNAs including ribosomal RNA (rRNA), piwi interacting miRNA (piRNA), transfer RNA (tRNA), small interacting RNA (siRNA), small nucleolar RNA (snoRNA) and small nuclear RNA (snRNA) (**Table S1**). A total of 2092121 and 1983614 reads in replicate 1 and replicate 2 respectively, of HPW library were unmapped after the RFAM blast, while 3906027 and 6616015 reads were unmapped in replicate 1 and replicate 2 respectively, in LPW library (**Table 1**). Length distribution analysis of the finally filtered reads showed preferred occurrence of 24 nucleotide long reads (**Figure S2**) in the sequenced libraries. Correlation of replicates of HPW and LPW library were significantly correlated to each other with r-value of 0.93 and 0.82, respectively (**Figure S3**).

**Table 1.** Mapping statistics of High Proanthocyanidin Winged bean containing Line (HPW) and Low Proanthocyanidin Winged bean containing Line (LPW) of *P. tetragonolobus* along with their two biological replicates.

| Samples | Raw Reads |  | After Filtration<br>(Q Value > 30 && adaptor) |  | 16-35 nt<br>(length) |  | RFAM |  |
| --- | --- | --- | --- | --- | --- | --- | --- | --- |
|  | Total | Distinct | Total | Distinct | Total | Distinct | Mapped | Unmapped |
| HPW_R1 | 45374213 | 4291092 | 38756906 | 3352329 | 20613214 | 2613885 | 521764 | 2092121 |
| HPW_R2 | 40713622 | 3660218 | 33564423 | 2987992 | 15382959 | 2455551 | 471937 | 1983614 |
| LPW_R1 | 47526600 | 6212028 | 45188452 | 5399167 | 26313706 | 4579360 | 673333 | 3906027 |
| LPW_R2 | 66054417 | 10038596 | 61240301 | 8702929 | 37258688 | 7426258 | 810243 | 6616015 |

### Identification of conserved and differential expressed miRNAs

A total of 32 and 50 conserved miRNA families were identified in HPW and LPW lines of *P. tetragonolobus,* respectively. Thirty-one miRNA families were common to both the libraries and one being specific to HPW and 19 to LPW (**Figure 2A**). These miRNAs have significant higher expression in LPW than HPW with the *p*-value < 0.0001 (**Figure 2B**). The conserved miRNA families comprised of total 139 family members whose log2 expression value was visualized through the heatmap representation (**Figure S6**). Most of the expressed conserved miRNAs showed higher expression in LPW line while some miRNAs showed almost equal level of expression for instance mir2118, mi403, mir5255. All the identified miRNAs showed their conserveness in 52 different plant lineages (**Figure S4**). Of the identified miRNAs, most of the miRNAs were found to be present in *Glycine max* in both the HPW (1060 miRNAs) and LPW libraries (1101 miRNAs). This study confirmed that, *Glycine max* and *P. tetragonolobus* are closely related with each other. As compared to HPW sequenced library, LPW showed much higher number of conserved miRNAs that have been reported in Poaceae. (**Figure S4**).

**Fig. 2.**
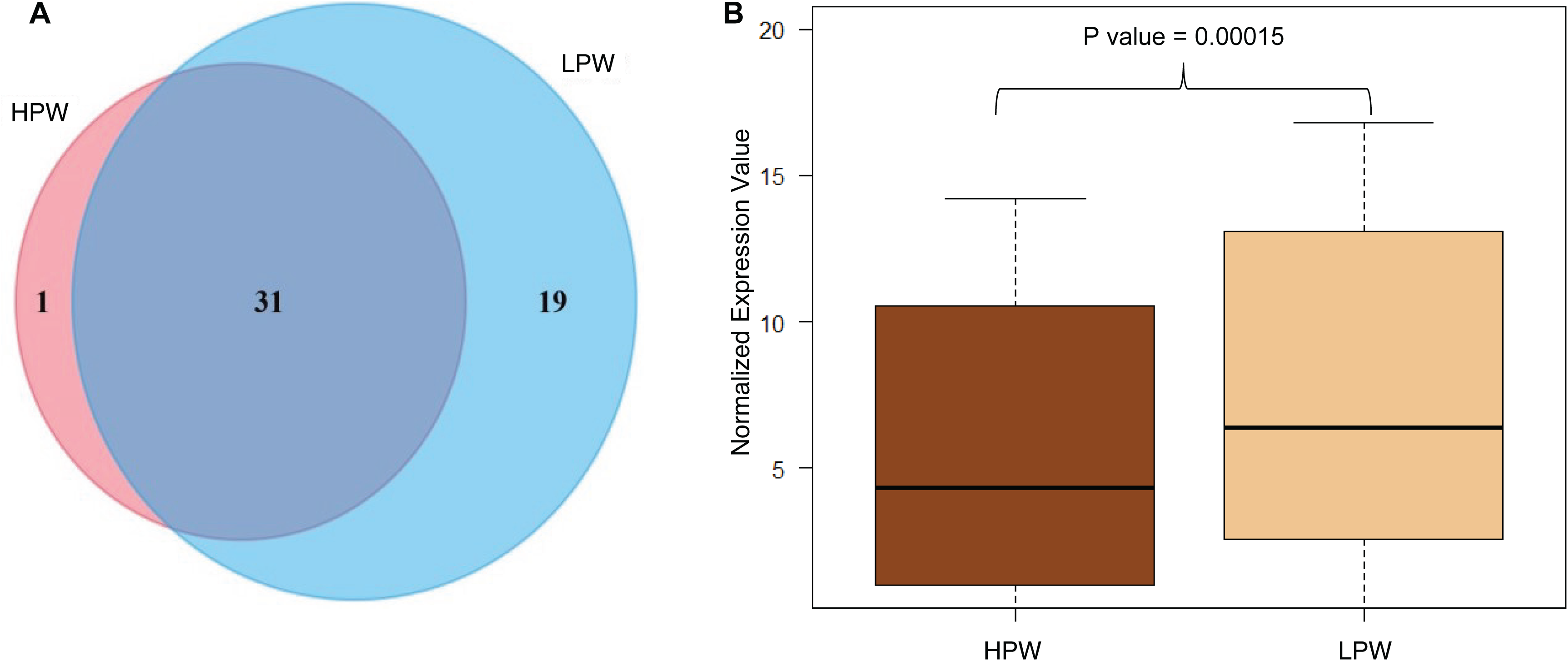
**A.** Identified conserved miRNAs in our sequenced library of two PA containing lines. Larger circle size represents the higher number of conserved miRNAs in LPW line while HPW has comparatively lower number of conserved miRNAs. **B.** Box plot representation of normalized expression value of conserved miRNAs in HPW and LPW lines. miRNAs expression level differences between two diverse lines are statistically significant (two sample t-test) with the p-Value 0.00015.

Among the identified conserved miRNAs, 23 miRNAs were differentially expressed between the two contrasting lines of *P. tetragonolobus* with log_2_ fc differences > ± 1, *p*-value < 0.005 and FDR value ≤ 0.05 (**Figure S5**). In HPW line, five miRNA families (i.e. mir319p, mir9726, mir862a-b and mir894) were significantly upregulated with fold change > 1 whereas, eight miRNA families (mir4416c-3p, mir396 (a-h, k), mir4414b) were significantly downregulated (**Table 2**).

**Table 2.**
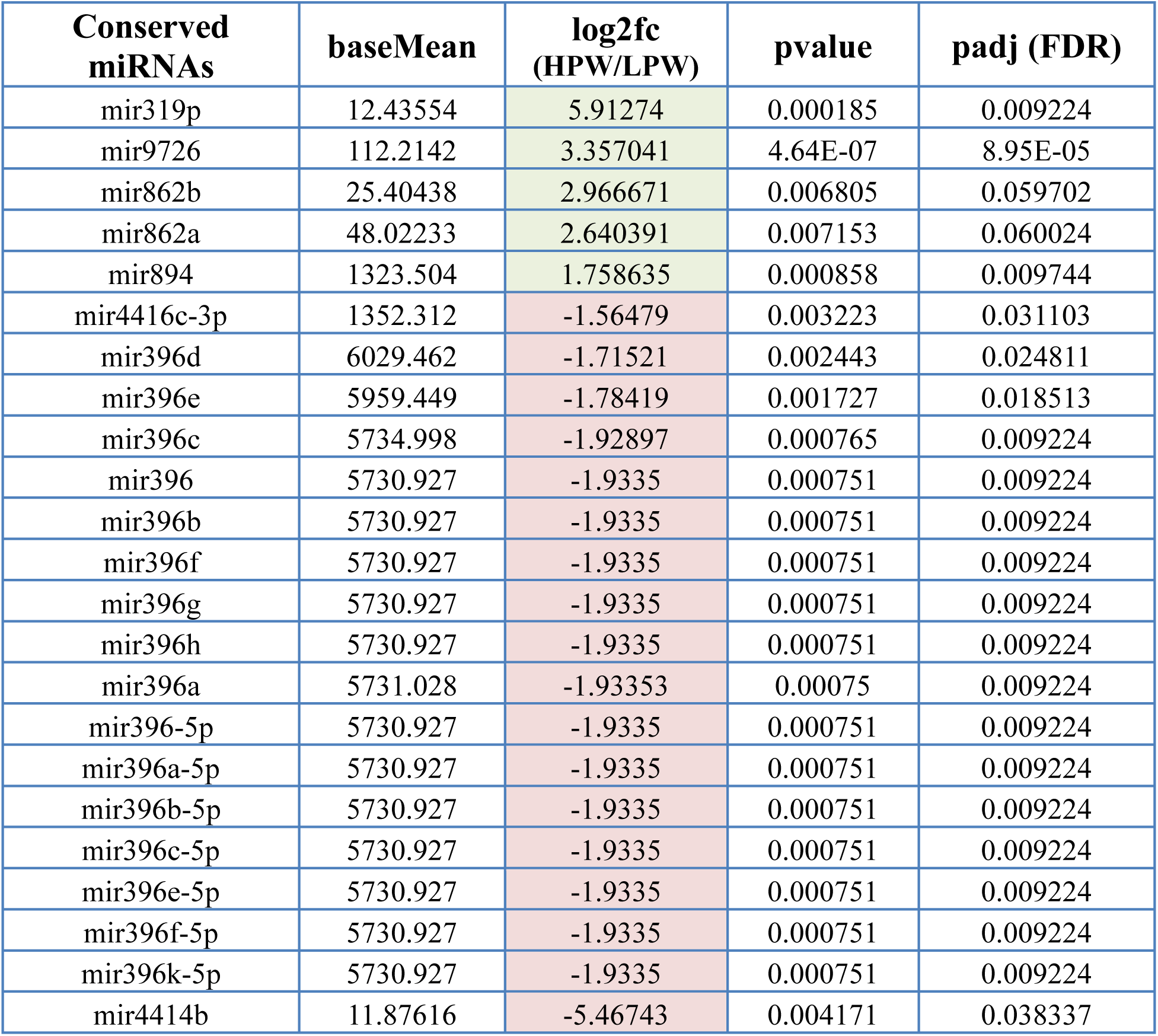
Mean expression value of differentially expressed miRNAs along with their log2fold change differences, pValue and FDR value. Green and red colored cell was used for upregulated and downregulated miRNAs in HPW line in compare to LPW line.

### Functional enrichment of miRNAs with their putative targets

Differentially expressed (DE) miRNAs in HPW and LPW lines of *P. tetragonolobus* were subjected to identification of their putative targets in *Glycine max* reference genome with seed size ranging from 2 to 13 nts, translation inhibition ranged from 10 to 11 nts and expectation score > 3. The differentially expressed miRNAs putatively targeted a total of 627 genes (3533 transcripts) **(Figure 3)** in which 555 genes were inhibited by the cleavage mechanisms while 72 genes were modified by the translation machinery (**Table S2**).

**Fig. 3.**
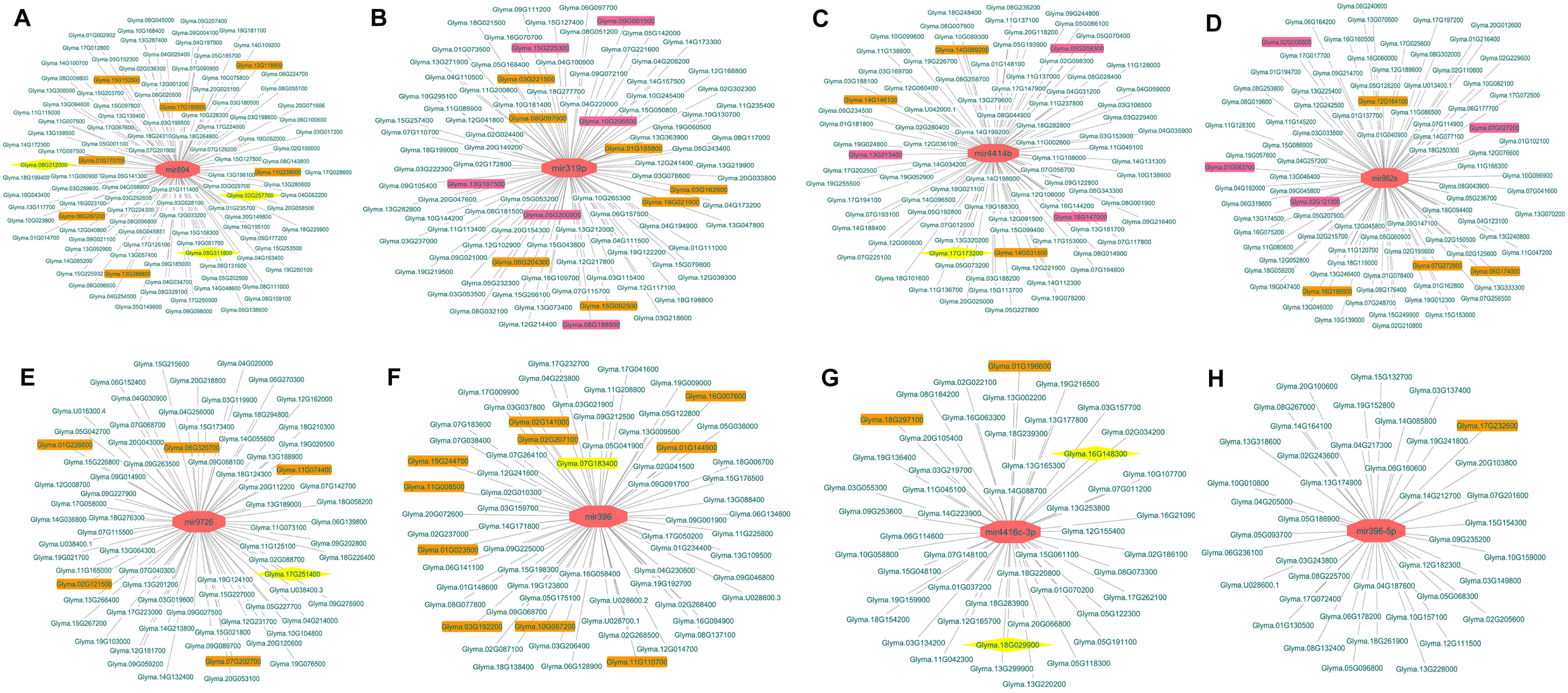
Target visualization of Differentially Expressed (DE) miRNAs in HPW and LPW lines of *P. tetragonolobus* for **A.** mir894 **B.** mir319p **C.** mir4414b **D.** mir862a **E.** mir9726 **F.** mir396 **G.** mir4416-c3p and **H.** mir396-5p. Diamond shaped pink color (node) represents the DE miRNAs and the targets were represented in edges with turquoise colored *Glycine max* Gene ID. Transcription Factors were displayed through the brown rectangular boxes, while secondary metabolites related genes were showed in pink colored boxes. The yellow colored edges are the validated targets of the corresponding miRNA (on the basis of literature).

The targeted genes comprised of many transcription factors (TFs) like GRF, MYB, bHLH, ARF, ERF, SBP TCP, C2H2 (**Figure 3**) (**Table S3**). These were targeted by cleavage mode of action. Exceptionally, mir894 inhibits Glyma.13288800 (HD-ZIP) and mir319p inhibits Glyma.08G188900 (MYB) TFs by translation activity. Secondary metabolite biosynthesis related genes were also inhibited by the differentially expressed miRNAs (**Figure 3**) (**Table S4**) in which mir4414b targets to dihydroflavonol-4-reductase (DFR, Glyma.17G173200), mir894 targeted to spermidine hydroxycinnamoyl transferase (SHT, Gyma.08311800, Glyma.08312000) and 2OG and Fe-dependent oxygenase (Gyma.02G257700). Mir4416c-3p targeted to SHT (Gyma.16G148300) and acyl transferase gene (Gyma.18G029900) while mir9726 targeted to oxidoreductases gene family (Gyma.17G251400).

Gene ontology analysis of putative targets suggested that they have functional role in negative regulation of growth (GO:0045926), histone modification (GO:0016570), floral organ formation (GO:0048449), Ca^2+^ transmembrane transport (GO:0070588) and thiamine pyrophosphate transport (GO:0030974) as well as in hydrolase activity (GO:0004553), maintaining the anion (GO:0055081) and amino acid homeostasis (GO:0080144) (**Figure 4A**). It also has a significant role in leaf morphogenesis (GO:0009965) and leaf development (GO:0048366).

**Fig. 4.**
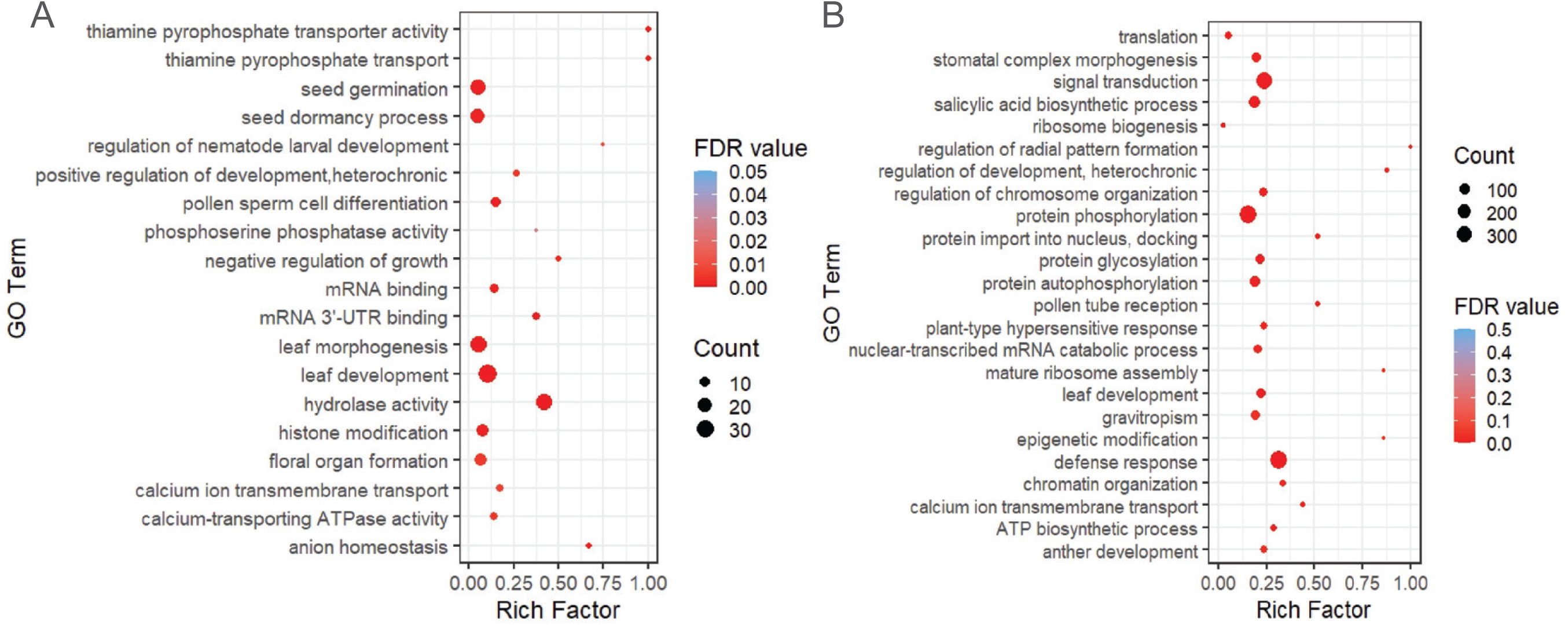
Gene ontology enrichment analyses of predicted targets of **A.** Differentially expressed conserved miRNAs and **B.** Novel miRNAs. The X-axis represents the rich factor (total number of genes/background genes) and Y-axis depicted the different GO Term. Dots were used for representing the total number of genes present in input datasets and color code represents their significant enrichment. All selected GO term are significant with the FDR Value less than 0.005.

### Elucidation of novel miRNA in HPW and LPW lines of *P. tetragonolobus*

Identification of novel miRNAs were carried out by unmapped reads that were not assigned as known and conserved miRNAs in both the lines. A total of 3543844 and 9175080 reads of HPW and LPW lines respectively were mapped against the *Glycine max* reference genome. Approximately, 8.658% and 7.571% reads were uniquely mapped in HPW and LPW lines, respectively. These mapped reads were further used for the identification of novel miRNAs through mirDeep2 pipeline with significant randfold *p*-value and at least three sequencing read depth (**Table 3**). MirDeep2 predicted total 19 novel miRNAs in *P.tetragonolobus* in which five novel miRNAs were predicted from the HPW line while 14 novel miRNAs from LPW line (**Figure 5 A,B**). These miRNAs were considered as putative novel miRNAs in *P.tetragonolobus.* Provisional IDs were provided to each putative novel miRNA by taking “pte-NmiR” as prefix and their putative targets were also predicted (**Table S5**). The gene ontology (GO) analysis identified the targets of *P.tetragonolobus* novel miRNAs. The GO analysis suggested their potential role in signal transduction, leaf development and regulation of chromosome organization along with the epigenetic modifications (**Figure 4B**).

**Fig. 5.**
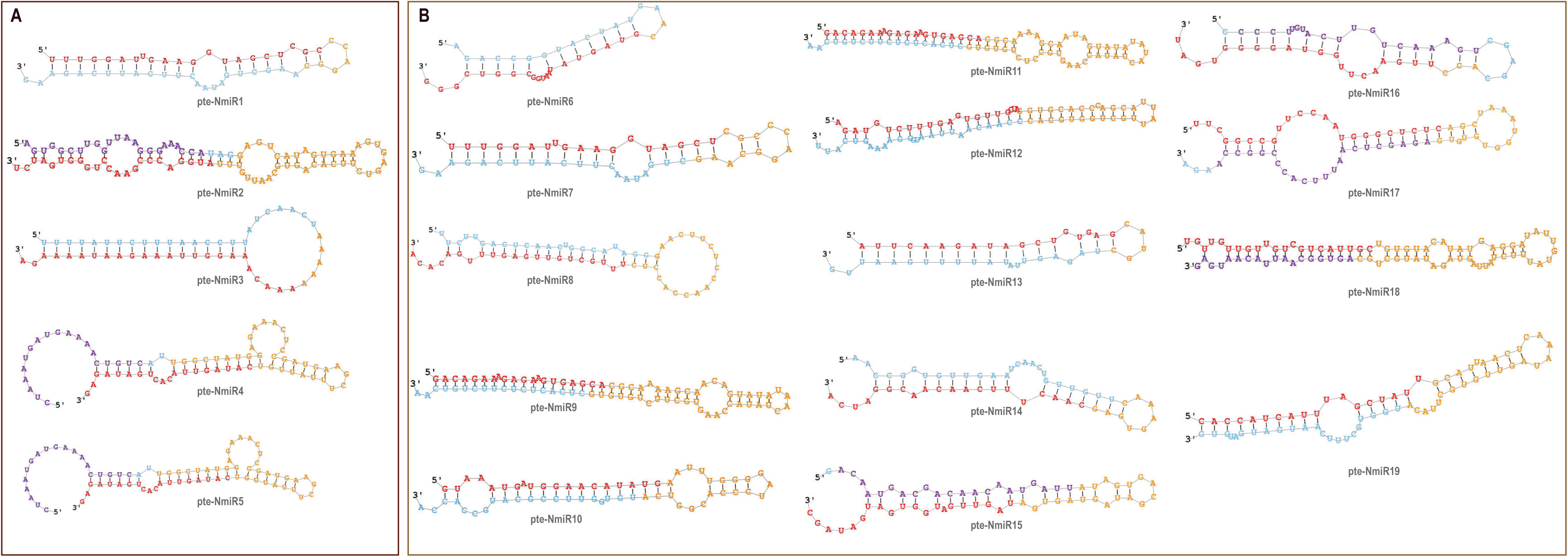
Secondary structure illustrations of predicted Novel miRNAs of *P. tetragonolobus* from the **A.** HPW and **B.** HPW sequenced Illumina library. Provisional Id was given to each predicted novel miRNAs in sequential manner with “pte-NmiR” prefixes. Star nucleotide sequences in miRNA secondary structures were represented in red in color.

**Table 3.**
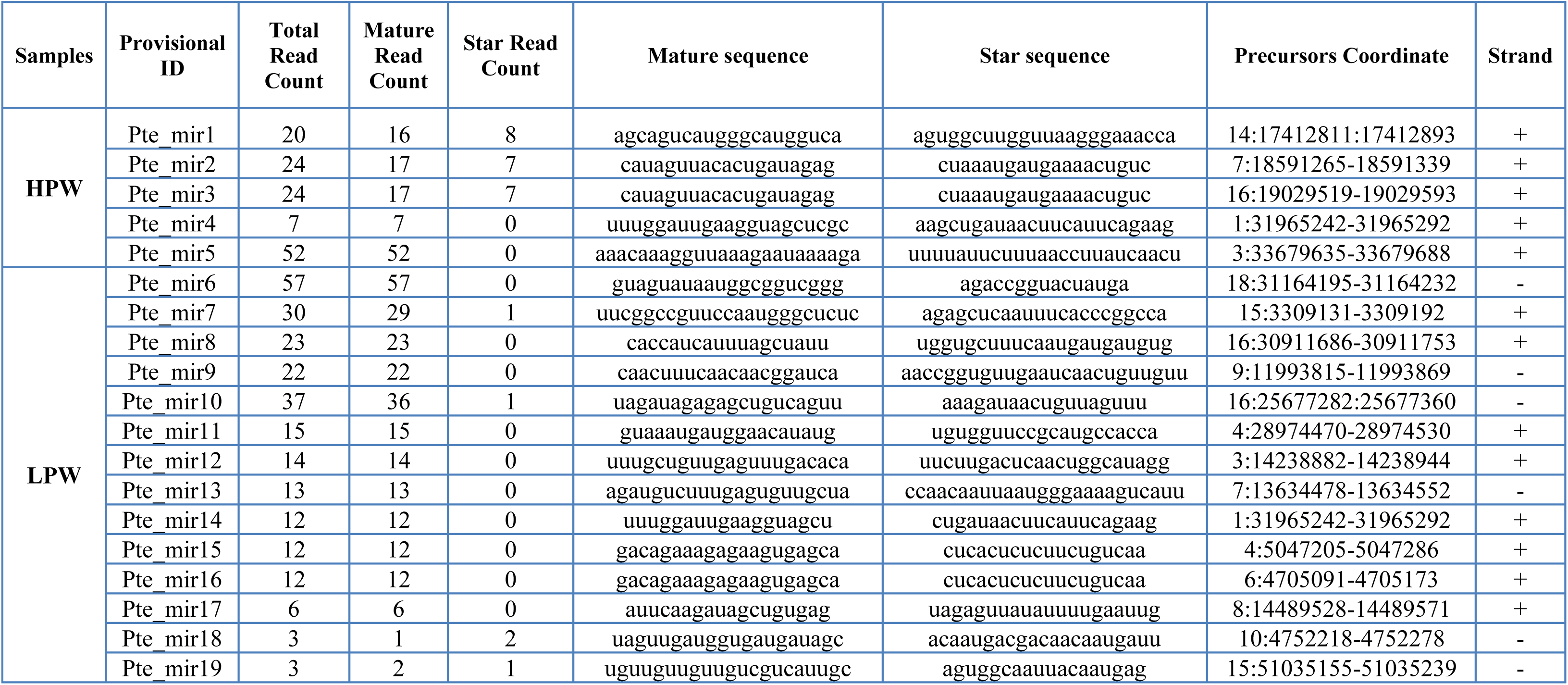
Detail information of predicted novel miRNAs in *P. tetragonolobus*. The provisional ID were given to all predicted novel miRNAs by adding “pte-NmiR” prefixes. Precursors coordinate of each miRNAs were mentioned along with their orientation on the genomic strand.

Apart from the novel miRNA predication, isoforms of the conserved miRNAs were also identified. These isoforms were varied in nucleotide length (1 to 7 nts) at either 3’ or 5’ overhangs with some nucleotide variation. (**Table 4**). These miRNA isoforms were named with “pte-IsomiR” prefixes along with their corresponding aligned miRNA ID.

**Table 4.**
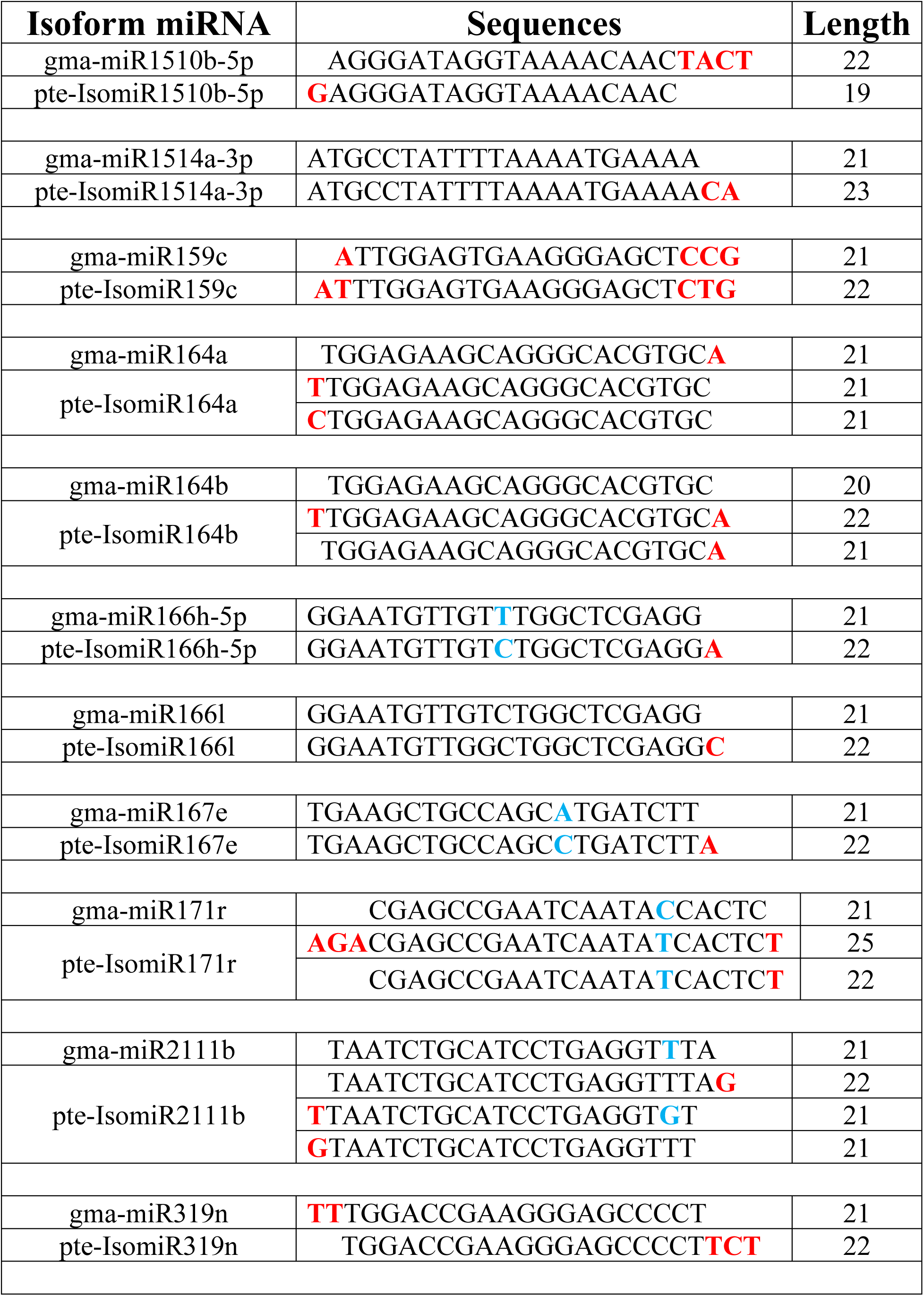
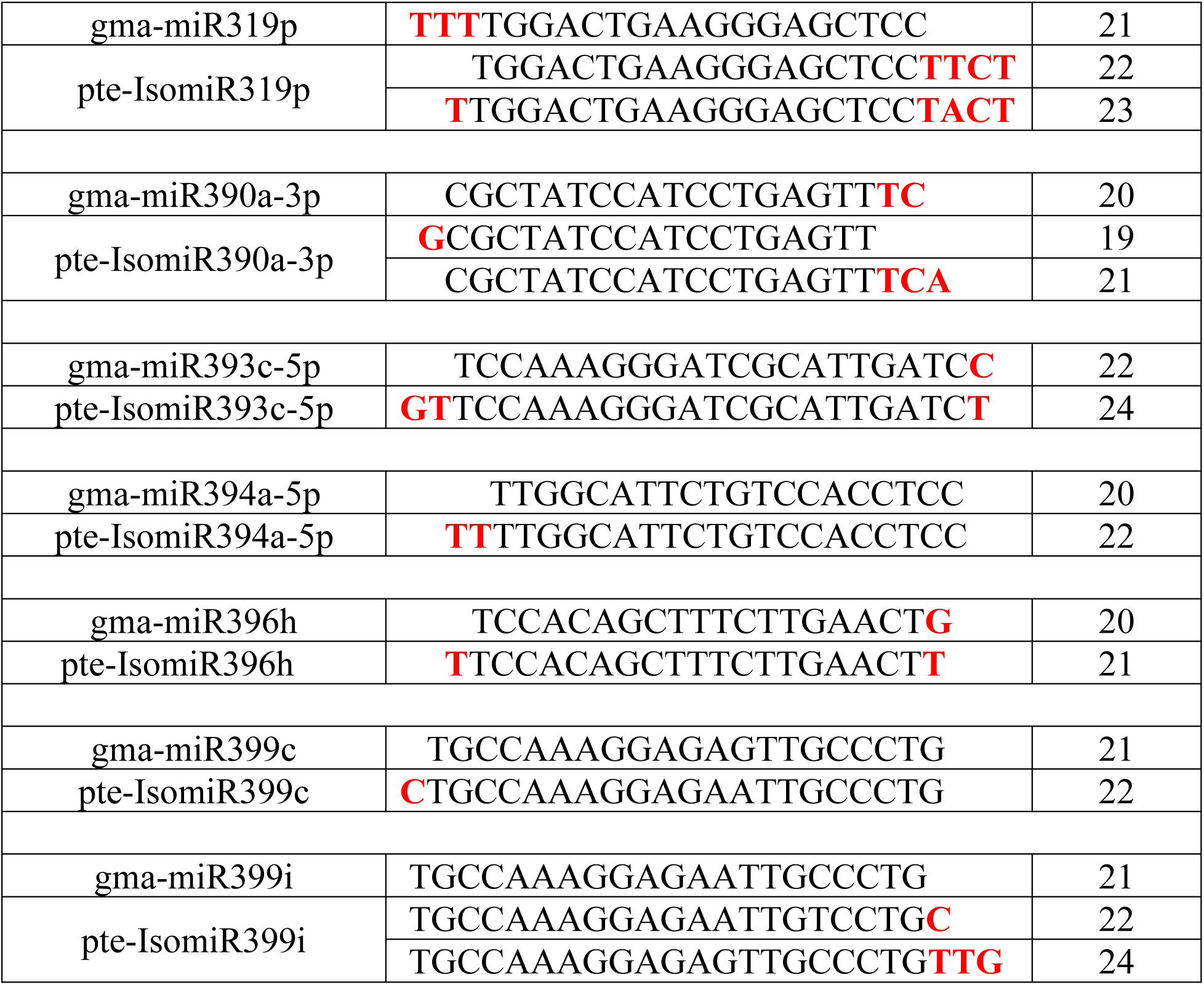
Identified miRNAs isoforms of *P. tetragonolobus* on the basis of *Glycine max* reference genome. All isoforms showed the nucleotides (nt) overhangs either at 5’ or 3’ marked with the red bold font letter. Blue font letter was used to displayed some nucleotide variation in the miRNA sequences.

### Real time expression analysis of the miRNAs in *P.tetragonolobus*

Validation of the expressed miRNAs was carried out between the contrasting lines (HPW and LPW) of *P.tetragonolobus* using stem-loop qRT-PCR. Twelve miRNA families including two novel miRNAs were randomly selected for validation. Out of the selected miRNAs, ten miRNAs were validated through qRT-PCR expression analysis. The qRT-PCR based miRNA expression analysis correlated with the sequenced libraries wherein LPW line showed higher number of log_2_ fold change differences in comparison to HPW line of *P. tetragonolobus* (**Figure 6**), with exception of miR862b which showed contrasting qPCR results vis-à-vis deep sequencing result. The two novel miRNAs exhibited same expression as in the sequencing result between the contrasting lines. Among the known miRNAs mir4416c-5p, miR2111j, miR156t, miR396, miR4414b, miR894, miR5139 were found to be differentially expressed and validated on qPCR platform. However, MiR862 exhibited to express at higher level in HPW line in deep complementing to its higher expression in LPW as validated through qPCR-based analysis. The primers used for qPCR analysis are listed in supplementary file (**Table S6)**.

**Fig. 6.**
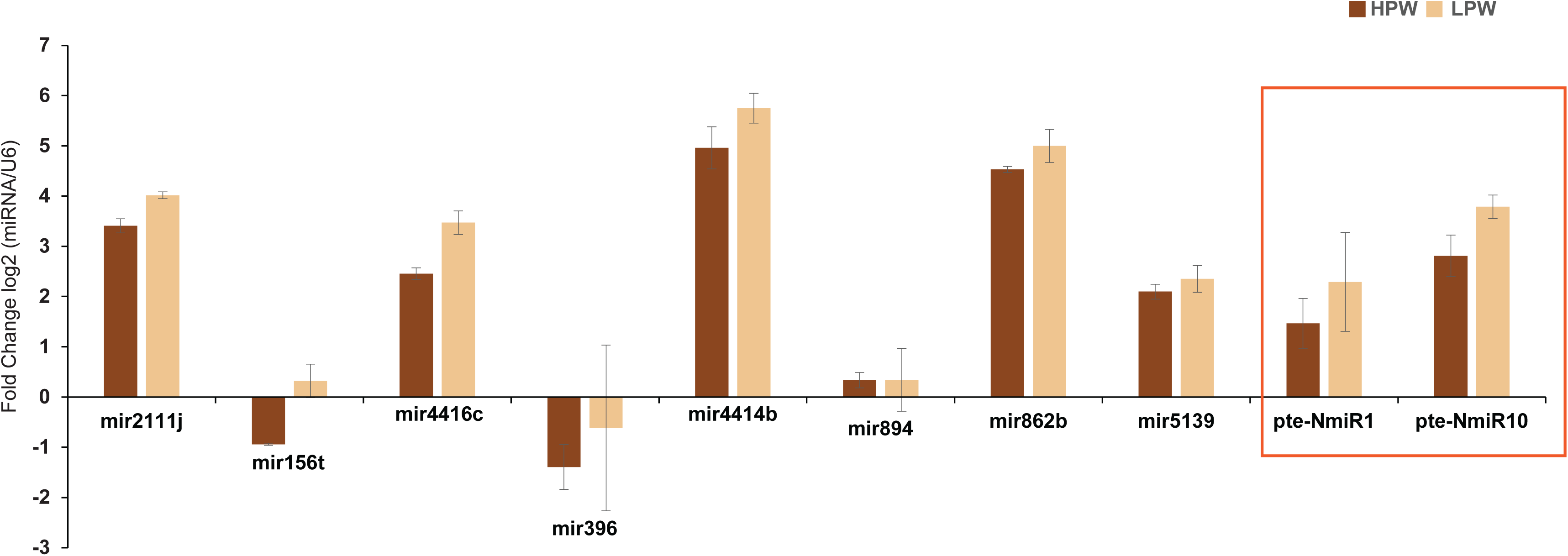
Real time Expression analyses of conserved and novel miRNAs in two contrasting lines (HPW and LPW) of *P.tetragonolobus*. The relative expression levels of the selected miRNAs were calculated using the 2–ΔΔCT method. The U6 gene was used as a Control. Each experiment consisted of three replicates and the error bars represent the standard deviation of the mean expression values among the replicates. Novel miRNAs were enclosed in red box.

## Discussion

MiRNAs have emerged as the key regulators of numerous biological processes including secondary metabolite synthesis and their distribution. Among the secondary metabolites, flavonoids are the important group of secondary metabolites in plants. They provide color and resistance against various pathogen attack to the plants. The flavonoid biosynthesis has been widely studied in many plants and the pathway operates in almost every fruit and vegetables and thus become a part of our regular diet (Tohge et al., 2017). Proanthocyanidins belong to flavonoid class of secondary metabolites and their biosynthesis and genetic regulation mechanism is yet to get fully explored. There are no reports till date regarding the miRNAs responsible for the biosynthesis of proanthocyanidins in *P. tetragonolobus*. This is the first attempt to report the responsible miRNAs for biosynthesis of proanthocyanidin in this underutilized legume *P. tetragonolobus*. Understanding the molecular basis of PA biosynthesis would help to manipulate its biosynthesis in legumes. This may pave way for altering the biosynthesis of PA.

The presence of different monomeric subunits of PA (catechin and epigallocatechin gallate) in varying levels in HPW and LPW lines of *P. tetragonolobus* suggests that, the PA level is possibly being regulated at different genetic levels. As leaf is one of the primary sites of secondary metabolite biosynthesis, so the miRNA analysis of leaves will be helpful for understanding the mechanism of PA biosynthesis and its further regulation. MiRNAs have already been identified to take part in various developmental processes including secondary metabolite synthesis e.g., miR858 & miR156 in flavonoid synthesis pathway and miRNA & miR414 & miR1134 in terpenoid biosynthesis(Gupta et al., 2017). High-throughput sequencing has enabled identification and deposition of miRNAs in miRbase. This enables database the process of identifying miRNAs in new plant species has become more accurate (Sripathi et al., 2018).

In the present study, approximately 50 million Illumina reads were sequenced and analyzed. This leads to the identification of 32 and 50 miRNA families in HPW and LPW lines respectively. The size distribution of the filtered reads showed the occurrence of 24nt small RNAs in both the lines. This result is consistent with the previously reported sRNA data in *Asparagus officinalis* (Chen et al., 2016), *Medicago truncatula* (Szittya et al., 2008) and *Citrus trifoliata* (Song et al., 2010). The fully sequenced and annotated *Glycine max* genome enabled to identify the conserved and novel miRNAs with their putative targets in *P. tetragonolobus*. *G. max* is a model legume and showed higher number of conserved miRNAs in the generated sequenced data of both the lines of *P. tetragonolobus*. The miRNAs identified through homology-based methods had revealed some highly conserved and non-conserved miRNA families in wide range of plants. The highly-conserved miRNA families include miR156, miR396, miR157, miR319 (Zhang et al., 2006) that are involved in many vital biological processes of plant growth and development. MiR4414, miR4416, miR5037, miR2111, miR9726 and miR894 are the miRNA families whose functions are yet to be elucidated. MiR156, miR396, miR4414b, miR408 were reported to be differentially expressed in HPW and LPW lines and some novel miRNAs were too found to be differentially expressed in the contrasting lines of *P. tetragonolobus*.

Most of the miRNA families were found to be highly expressed in LPW than HPW lines of *P. tetragonolobus*. Out of the differentially expressed miRNAs, miR4414b, miR4414c, miR396, miR156, and miR894 have some direct or indirect control over proanthocyanidin biosynthesis (Gou et al., 2011; Gupta et al., 2017; Wang et al., 2020). MiR156 interacts with *SQUAMOSA PROMOTER BINDING PROTEIN-LIKE (SPL)* gene to increase the levels of anthocyanins and regulate the levels of other associated products like flavones and flavanols. *SPL* genes have been found to be negative regulators of flavonoid biosynthetic pathway as they disrupt MYB-bHLH-WD40 ternary complex which act as activators of late biosynthetic genes. Thus, SPL9 affects the accumulation of anthocyanin and downstream compounds (Gou et al., 2011; Wang et al., 2020). Differential expression of miR156 suggests its possible role in the regulation of proanthocyanidin synthesis in *P. tetragonolobus*. Dihydroflavonol reductase (*DFR*) gene was also found to be targeted by the miR4414b, which was found to be differentially expressed and also has been validated. DFR is one of the key regulatory-enzymes of the flavonoid biosynthetic pathway that is essential for proanthocyanidin synthesis and accumulation (Li et al., 2017). Higher level of expression of miR441b in LPW line suggests its possible role in lowering the PA level in LPW leaves. Higher expression of differentially expressed miR396 negatively regulates flavonoid synthesis targeting the *GRF8* gene, a positive regulator of flavanone-3-hydroxylase (*F3H*) gene (Dai et al., 2019) of the pathway. Moreover, target prediction shows UDP-glucosyl transferase which glycosylates anthocyanidin is targeted by miR396 and the glycosylation step usually leads to the production of various anthocyanin pigments (Zhao et al., 2012) and the unused anthocyanidins are directed for preptechin (monomeric unit of PA) formation by the activity of anthocyanidin reductase enzyme (He et al., 2008). UDP-glycosyl transferase is a large gene family having different roles; one being involved in secondary metabolite synthesis. UGT78D2 that was predicted to be one of the targets of miR396 encodes for flavonoid 3-O-glucosyltransferase, this catalyzes the glucosylation of both flavonols (quercetin) (Kim et al., 2012) and anthocyanidins at the 3-OH position (Pourcel et al., 2010). Moreover, overexpression of UDP-glycosyl transferase gene produces higher amounts of anthocyanin and proanthocyanidin (Rao et al., 2019). Differential expression of miR396 may be correlated with the PA synthesis. MiR396 is functionally validated with its predicted target i.e., UDP-glucosyl transferase (*UGFT*) gene, the foresaid hypothesis can be a major lead in this study. Other miRNAs which have been identified with their targets playing role in PA synthesis are miR4414b, miR4414c and miR894. They have differential expression pattern in HPW and LPW lines of *P. tetragonolobus,* suggesting their probable role in PA biosynthesis. A putative model has been illustrated from the high throughput sequencing analysis of two contrasting lines to highlight the probable role of miRNAs in proanthocyanidin biosynthesis in *P. tetragonolobus* (**Figure 7**).

**Fig. 7.**
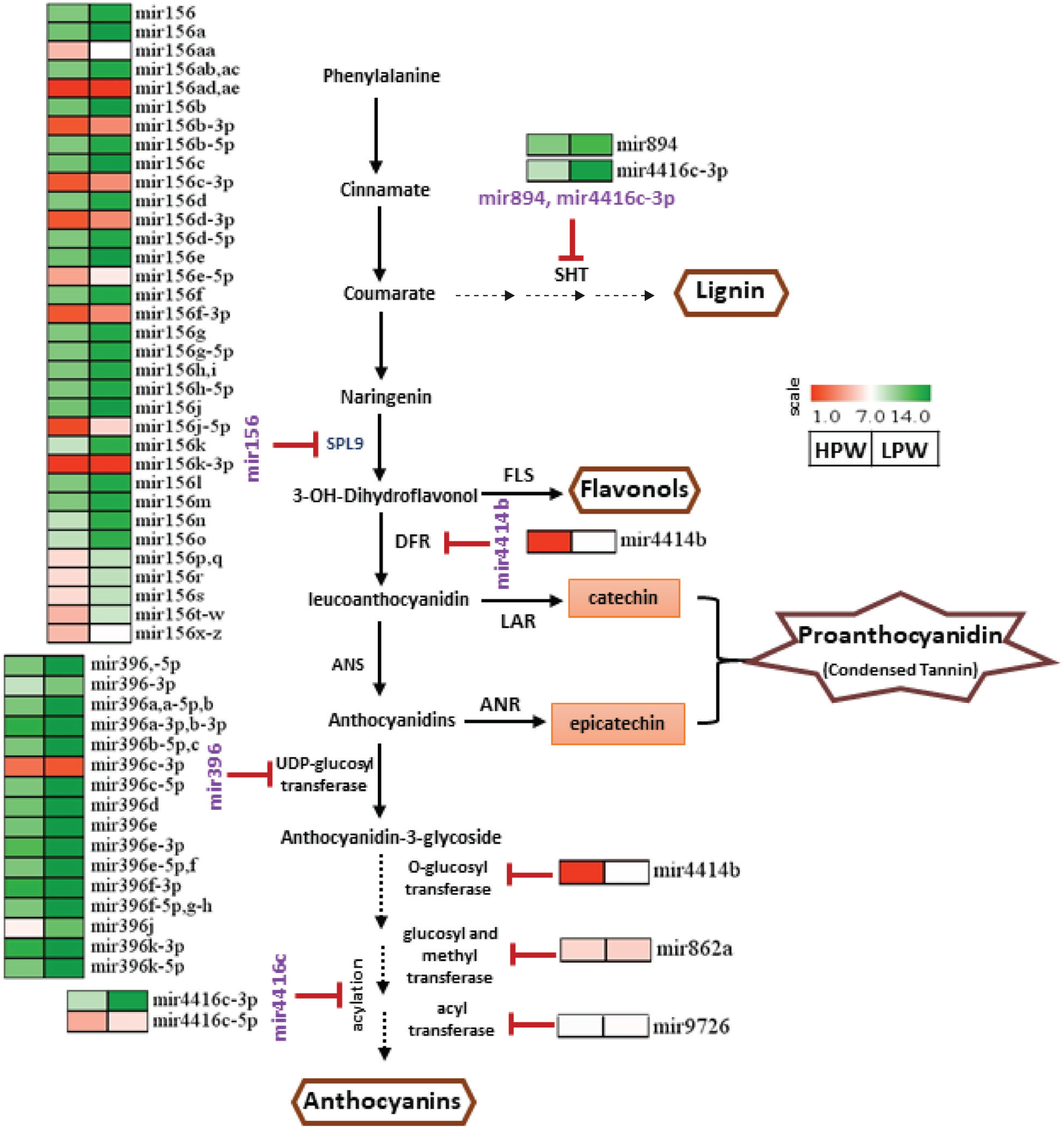
A putative model that illustrates the role of miRNAs in different proanthocyanidin content in the two contrasting line of *P. tetragonolobus*. Continuous arrow lines displayed the known pathways while break arrow showed the some missing pathway link. Red stop lines indicate the putative targets of the expressed miRNAs in the two sequenced library. The expressed miRNAs that hindering the flavonoid pathway are written in purple font color and their expression value was represented through the heatmap with the coloring scale ranging from 1 to 14. Left side and Right side expression bar was used for HPW and LPW library, respectively.

## Conclusion

Two different lines of *P.tetragonolobus* were scrutinized though biochemical assays and termed as HPW and LPW lines. Various genes, transcription factors and enzymes related to the subsequent pathways of PA biogenesis were targeted by expressed miRNAs in the contrasting PA-containing lines. HPW line has low expression value of most of the miRNAs both in deep sequencing and qPCR. Flavonoid synthesis pathway is a complex network of sequential actions of enzymes and transcription factors. This leads to the synthesis of many important compounds (Vogt, 2010) including proanthocyanidin which is the focus of this study. miRNAs have emerged as the key modulators of gene expression and have been found to play a role in this pathway. In our study apart from the previously characterized miRNAs, certain conserved and non-conserved and novel miRNAs with potential targets in the pathway have been identified, indicating the clue for the molecular mechanism of PA metabolism in *P. tetragonolobus*. However, further experimental validation of the miRNAs and their targets can reveal the exact mechanism through which miRNAs take part in PA synthesis in *P.tetragonolobus*. Most of the miRNAs in the proanthocyanidin biosynthesis were upregulated in LPW library, which supports the higher level of proanthocyanidin production in HPW as compared to LPW. This work provides an information regarding the known and novel *Psophocarpus tetragonolobus* miRNAs, their targets and their possible role in PA metabolism. Further, *in vitro* characterization and validation of the identified conserved and novel miRNAs might provide an important clue to regulation of PA synthesis and accumulation in winged bean.

## Acknowledgement

CSM and SPN acknowledge Dept. of Biotechnology for the financial support to carry out this activity and Director, CSIR-NBRI for providing the infrastructure facility. We also acknowledge NBPGR for providing required germplasm. PP and SPN acknowledge University Grants Commission for research fellowship grant. Authors also acknowledge Kirti Pandey and Arpit Chauhan for providing the experimental support during the study.

## Author’s contribution

CSM and SPN designed the research work. SPN conducted most of the experiments and prepared sample for small RNA sequencing. PP performed all the bioinformatic analyses. SPN and PP wrote the manuscript. VS and AMT provided necessary suggestions in conducting experiments. CSM, VS and SB helped in finalizing the manuscript and critically assessed the report. All authors read and approved the manuscript.

## Figure Legends

**Fig. S1.**
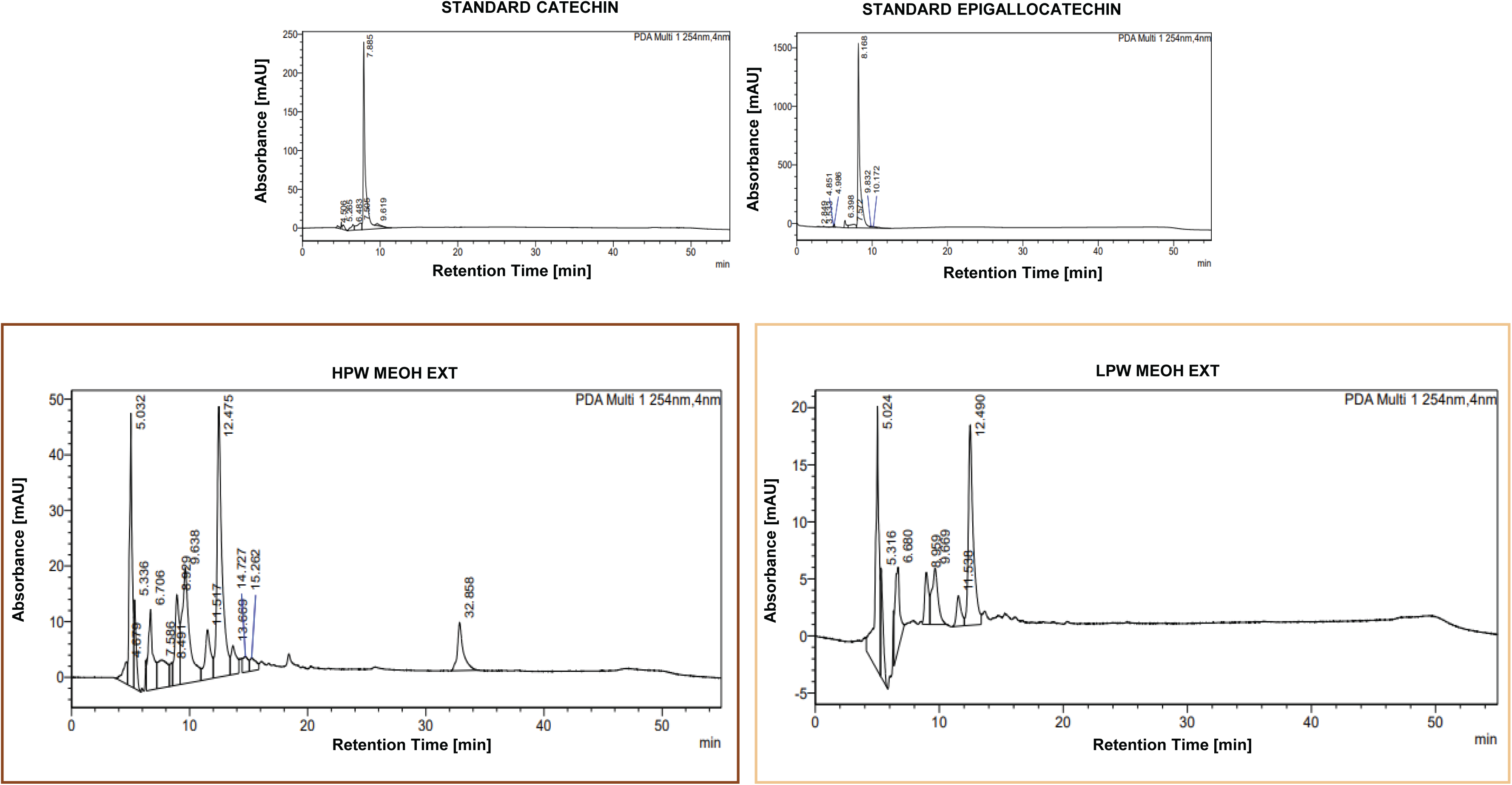
HPLC Chromatogram of standards (catechin and epigallocatechin gallate) and methanol extract of *P. tetragonolobus* leaves

**Fig. S2.**
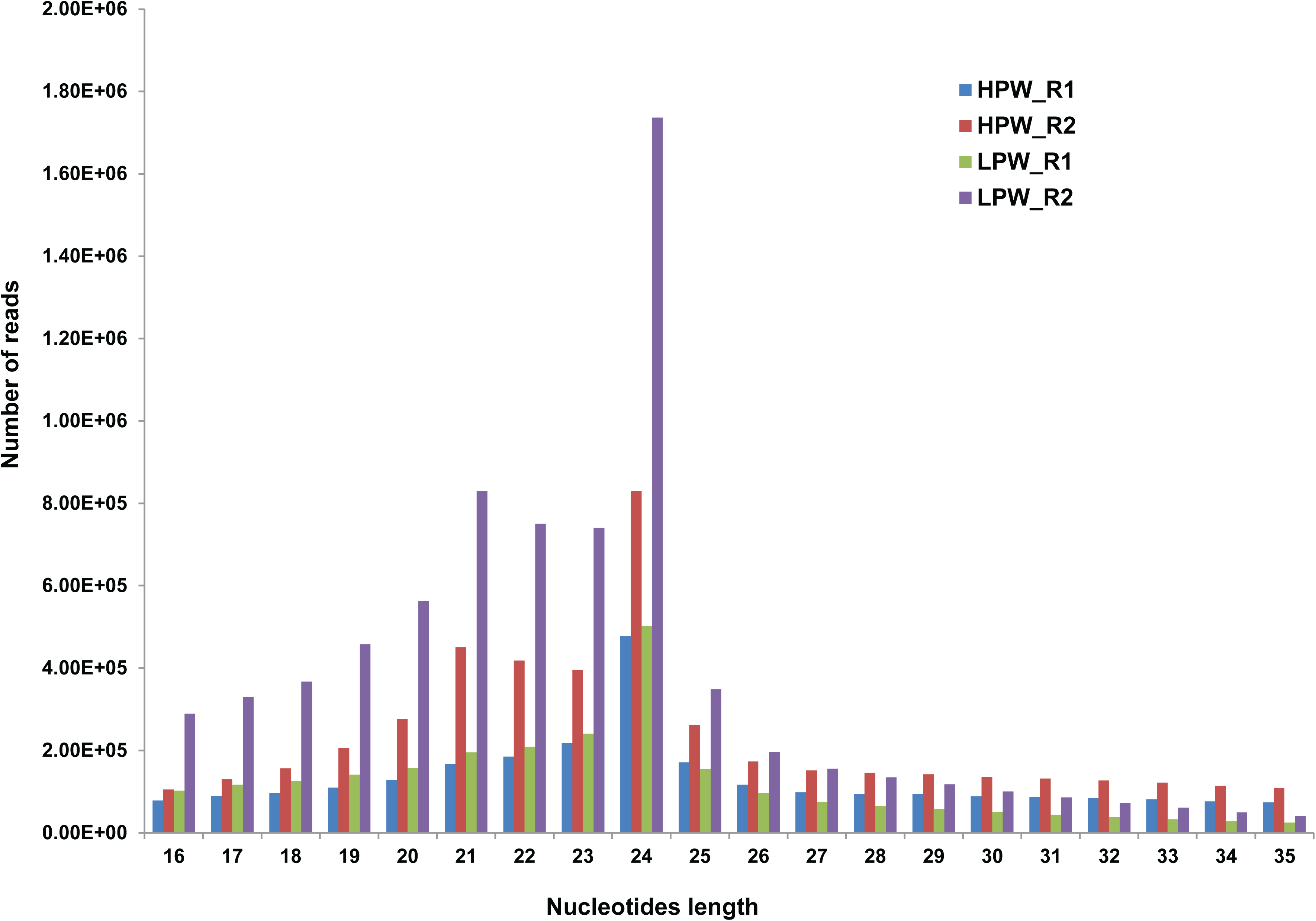
Nucleotides variation length of small mRNA in the sequenced library of HPW and LPW of *P. tetragonolobus*.

**Fig. S3.**
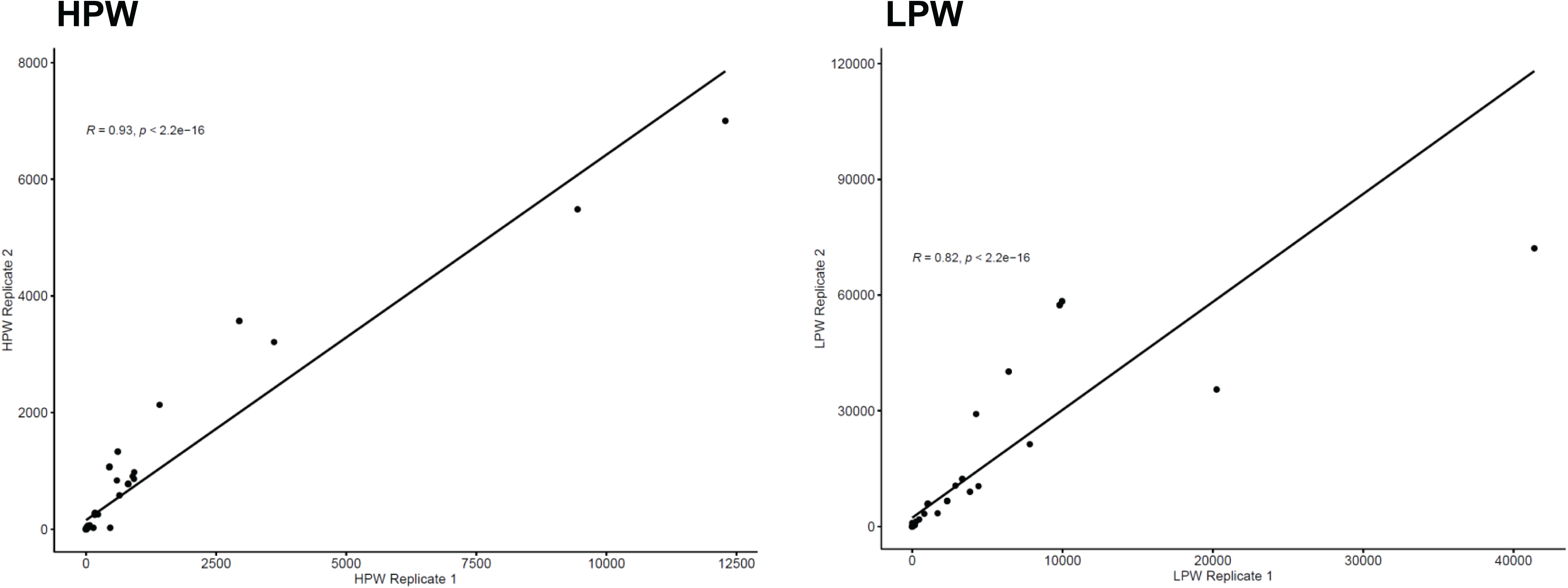
Correlation plot between the replicate 1 and replicate 2 of Illumina sequenced HPW and LPW library. R value represents the correlation value of 0.93 and 0.82 in HPW and LPW library, respectively with the significant p value < 2.2e-16.

**Fig. S4.**
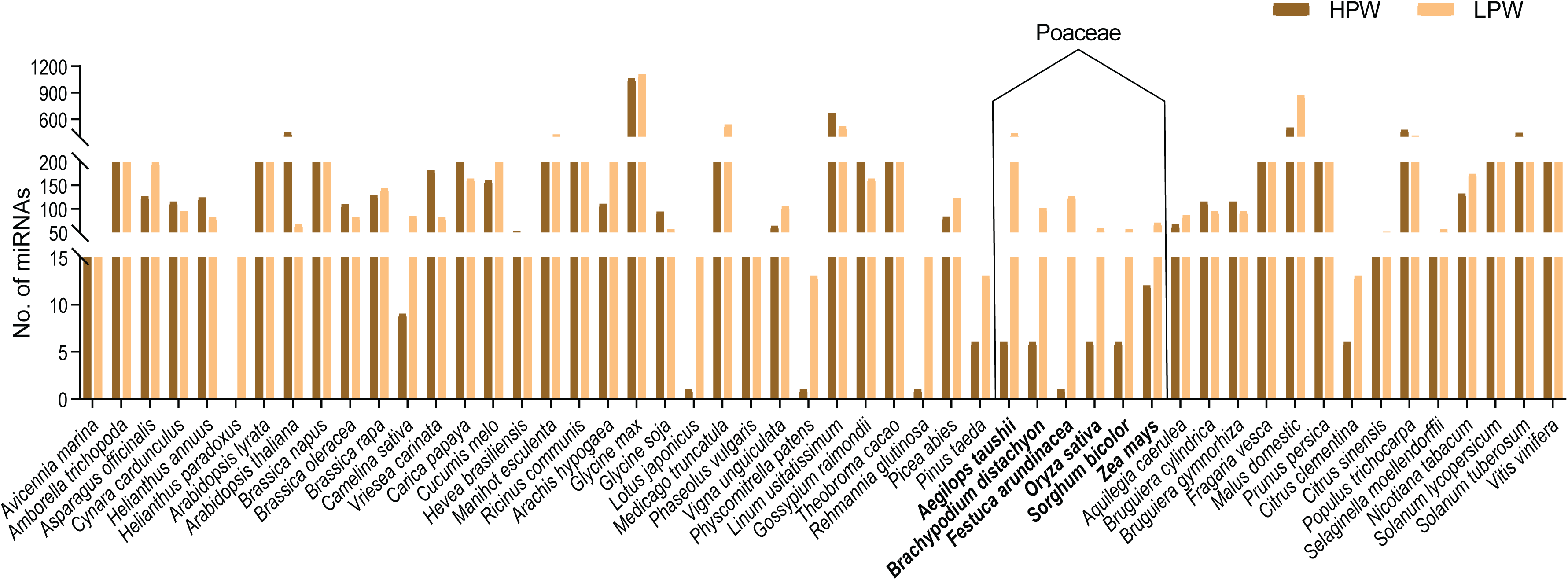
Different number of identified conserved miRNAs in HPW and LPW library in 52 plant lineages. X-axis represented the different plant species while different numbers of miRNAs was scaled in Y-axis. Bold font scientific name was used for plant species that belongs to Poaceae family.

**Fig. S5.**
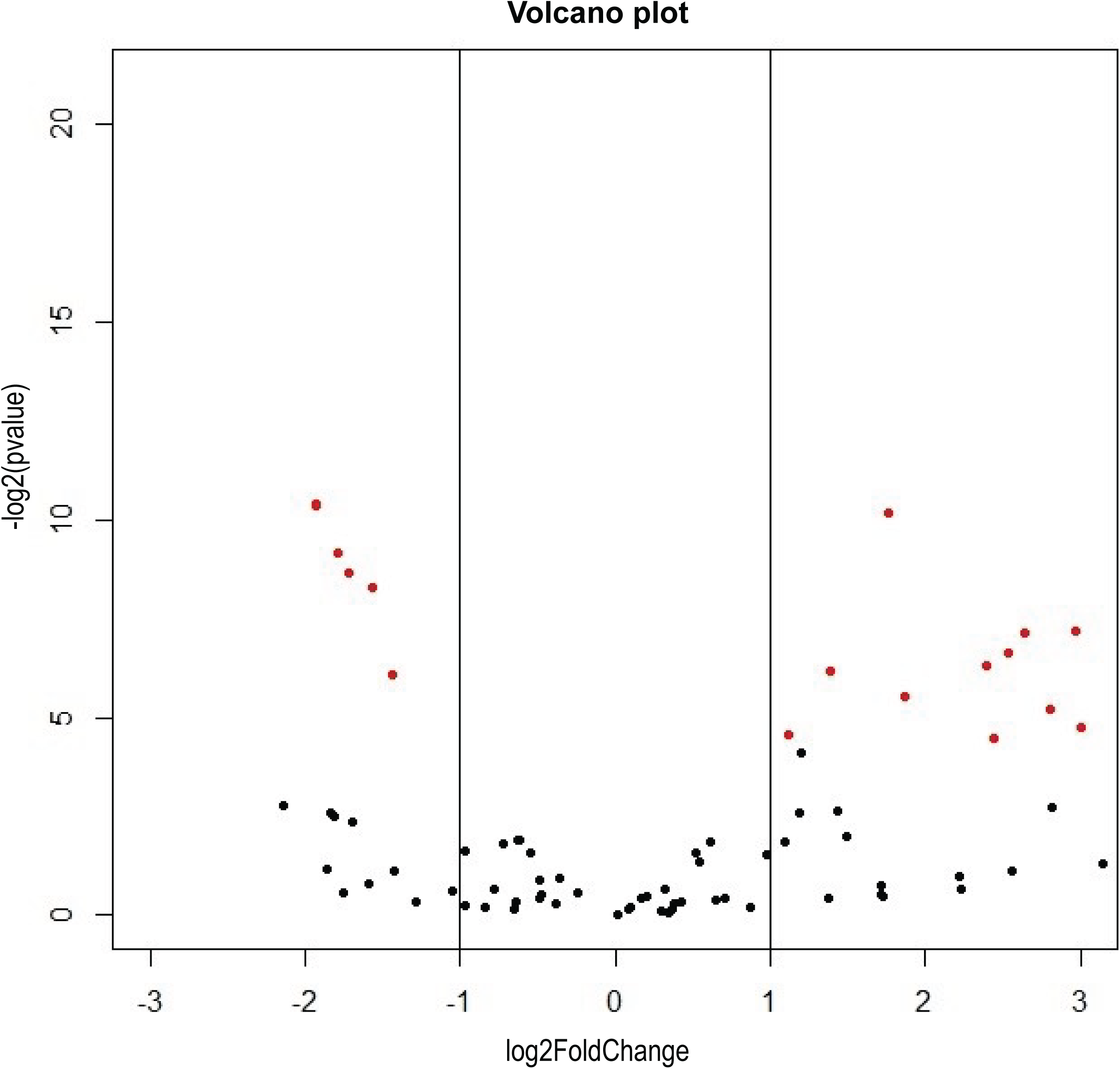
Volcano plot to represents the differentially expressed miRNAs bwteeen the HPW and LPW lines of *P. tetragonolobus*. X-axis was used for the log2Fold-change whereas Y-axis representing the –log2(pValue). Each dots describe the expressed miRNAs, wherein red dots define the differentially expressed miRNAs.

**Fig. S6.**
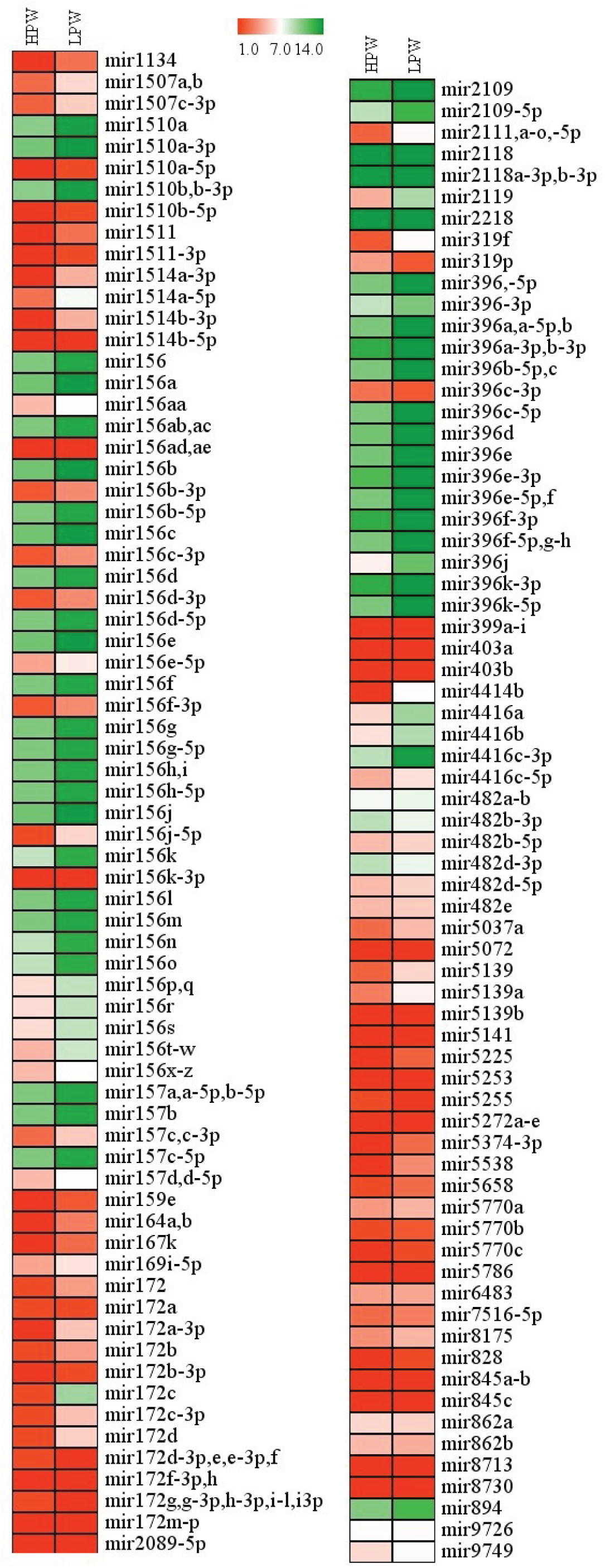
Heat map visualizations of log2fold change expression value of miRNAs from the High Proanthocyanidin Winged bean containing Line and Low Proanthocyanidin Winged bean containing Line. Lower expression value was represented in the red in color while higher expression was illustrated with green color. The color scale was ranged from 1 to 14.

